## Supplementary information for "A Recurrent Neural Circuit Mechanism of Temporal-scaling Equivariant Representation"

**5 Supplementary Figures** **10**

*Corresponding author.

Preprint. Under review.

### 1 Math background of the temporal scaling group

#### 1.1 Temporal scaling group and its generator

Several properties of the $\hat{S}(\alpha)$ can be directly derived from the definition (Eq. 2) in the main text [1],

$$\hat{S}(1) = 1\ , \tag{S1a}$$

$$\hat{S}(\alpha) \cdot \hat{S}(\beta) = \hat{S}(\beta) \cdot \hat{S}(\alpha) = \hat{S}(\alpha\beta), \tag{S1b}$$

$$\hat{S}(\alpha)^{-1} = \hat{S}(\alpha^{-1}). \tag{S1c}$$

Eq. (S1a) shows that a scaling transformation with factor 1 will leave $\mathbf{y}(t)$ unchanged. Eq. (S1b) indicates that sequential actions of two scaling operators can be composed into another scaling operator (start timing alignment is needed, see SI. Sec. 1.3), which ensures the *closure* of the group. Moreover, the composition is irrelevant with the order of the two operators, suggesting the temporal scaling group is *commutative*. At last, Eq. (S1c) shows the inverse of a temporal scaling operator is the one with an inverse scaling factor. According to the Lie group theory, generators can be derived by evaluating the derivative of the group element at the identity element. As for the temporal scaling group we are studying in the present work, we have:

$$\begin{aligned}
\hat{g} \cdot \mathbf{y}(t) &= \frac{d\hat{S}(\alpha)}{d\alpha}\Big|_{\alpha=1} \cdot \mathbf{y}(t) \\
&= \frac{\hat{S}(\alpha+\delta\alpha) - \hat{S}(\alpha)}{\delta\alpha}\Big|_{\alpha=1} \cdot \mathbf{y}(t) \\
&\approx \frac{[\mathbf{y}(\alpha t) + \delta\alpha t \partial_t \cdot \mathbf{y}(\alpha t) - \mathbf{y}(\alpha t)]_{\alpha=1}}{\delta\alpha} \\
&= t\partial_t \cdot \mathbf{y}(t).
\end{aligned} \tag{S2}$$

Therefore, we obtain that $\hat{g} = t\partial_t$. Here, $\mathbf{y}(t)$ denotes an arbitrary temporal sequence.

With the generator we have obtained, we can generate a set of continuous lie group elements by exponential mapping. Taking the derivative of $\hat{S}(\alpha)$ with respect to its parameter $\alpha$, and utilizing the property $\hat{S}(\alpha\beta) = \hat{S}(\alpha)\hat{S}(\beta)$, it yields:

$$\begin{aligned}
\frac{d\hat{S}(\alpha)}{d\alpha} &= \frac{\hat{S}(\alpha+\delta\alpha) - \hat{S}(\alpha)}{\delta\alpha} \\
&= \frac{\hat{S}(\alpha)\hat{S}(1+\delta\alpha/\alpha) - \hat{S}(\alpha)\hat{S}(1)}{\alpha \cdot \delta\alpha/\alpha} \\
&= \frac{\hat{S}(\alpha)}{\alpha} \cdot \frac{\hat{S}(1+\delta\alpha/\alpha) - \hat{S}(1)}{\delta\alpha/\alpha} \\
&= \frac{\hat{S}(\alpha)}{\alpha} \cdot \hat{g}.
\end{aligned} \tag{S3}$$

Solving this equation, we obtain the exponential map of the generator,

$$\hat{S}(\alpha) = \exp(\ln\alpha \cdot \hat{g}), \tag{S4}$$

then we have a temporal scaling operator with scaling factor $\alpha$. To verify the above exponential form of the TS operators, we apply it to $\mathbf{y}(t)$ and see whether it generates the same TS effect as defined in Eq. (2).

$$\begin{aligned}
\exp(\ln\alpha \cdot \hat{g}) \cdot \mathbf{y}(t) &= \lim_{\epsilon\to 0} \exp(\ln\alpha/\epsilon \cdot \epsilon\hat{g}) \cdot \mathbf{y}(t) \\
&= \lim_{\epsilon\to 0} (1+\epsilon\hat{g})^{\ln\alpha/\epsilon} \cdot \mathbf{y}(t) \\
&= \lim_{\epsilon\to 0} \mathbf{y}\left[(1+\epsilon)^{\ln\alpha/\epsilon} t\right] \\
&= \mathbf{y}[\exp(\ln\alpha)t] \\
&= \mathbf{y}(\alpha t),
\end{aligned} \tag{S5}$$

where we used $\lim_{\epsilon \to 0}(1+\epsilon)^{1/\epsilon} = e$. And from the 2nd to the 3rd row in the above equation, we apply the $\ln \alpha/\epsilon$ infinitesimal transformations $(1+\epsilon\hat{g})$ to $\mathbf{y}(t)$. Eq. (S5) verified that the exponential map (Eq. S4) is indeed an operator of the TS group.

The generator and its exponential map can also be derived from the definition. As stated in the main text, $\hat{S}(\alpha)\cdot\mathbf{y}(t) = \mathbf{y}(\alpha t)$. We can represent $\mathbf{y}(\alpha t)$ as $\mathbf{y}(\alpha t) = \mathbf{h}(\ln t + \ln \alpha)$, where $\mathbf{h}(x) = \mathbf{y}(e^x)$. Then we apply a full-order Taylor expansion to $\mathbf{h}(\ln t + \ln \alpha)$ as follows:

$$\mathbf{h}(\ln t + \ln \alpha) = \sum_{n=0}^{\infty} \frac{(\ln \alpha)^n}{n!} \mathbf{h}^{(n)}(\ln t), \tag{S6}$$

where $\mathbf{h}^{(n)}(\ln t)$ represents the n-th order derivative of $\mathbf{h}$ with respect to $\ln t$. We can calculate the first-order derivative as $\mathbf{h}^{(1)}(\ln t) = \frac{d\mathbf{y}(t)}{d\ln t} = t\frac{d\mathbf{y}(t)}{dt}$. Using this result, we can derive the n-th order derivative as:

$$\begin{aligned}
\mathbf{h}^{(1)}(\ln t) &= \frac{d\mathbf{y}(t)}{de^{\ln t}}\frac{de^{\ln t}}{d\ln t} = \frac{d\mathbf{y}(t)}{d\ln t} = t\frac{d\mathbf{y}(t)}{dt} \\
\mathbf{h}^{(2)}(\ln t) &= \frac{d}{d\ln t}\frac{d\mathbf{y}(t)}{d\ln t} = \left(t\frac{d}{dt}\right)^2 \mathbf{y}(t) \\
&\cdots \\
\mathbf{h}^{(n)}(\ln t) &= \left(t\frac{d}{dt}\right)^n \mathbf{y}(t),
\end{aligned} \tag{S7}$$

thus obtaining the expression of $\hat{S}(\alpha)\cdot\mathbf{y}(t)$ as:

$$\begin{aligned}
\hat{S}(\alpha)\cdot\mathbf{y}(t) = \mathbf{y}(\alpha t) &= \mathbf{h}(\ln t + \ln \alpha) \\
&= \sum_{n=0}^{\infty} \frac{(\ln \alpha)^n}{n!}\mathbf{h}^{(n)}(\ln t) \\
&= \sum_{n=0}^{\infty} \frac{(\ln \alpha \cdot t\frac{d}{dt})^n}{n!}\mathbf{y}(t) \\
&= \exp\left(\ln \alpha \cdot t\frac{d}{dt}\right)\cdot \mathbf{y}(t),
\end{aligned} \tag{S8}$$

which leads to the same results as Eq. S2 and S4.

##### 1.2 Temporal reversal operators

We note that the exponential form of TS operators (Eq. S4) is not consistent with temporal reversal operators ($\alpha < 0$), because a typical logarithmic function cannot take negative values. To resolve this notation issue, we effectively define a *generalized* logarithmic function that can take negative $\alpha$ as input. Specifically, we use the Euler's formula,

$$\exp(i\pi) = -1, \quad (i = \sqrt{-1})$$

Applying the "logarithm" on the LHS and RHS of the above equation,

$$\ln(-1) = i\pi,$$

we can get the notation of a generalized logarithmic function. For any negative number $\alpha$, we have,

$$\ln \alpha = \ln(-|\alpha|) = \ln(\mathrm{e}^{i\pi}|\alpha|) = i\pi + \ln|\alpha|.$$

Thus, the TS operators with negative factors $\alpha$ can be still denoted by using the same exponential map as defined in Eq. (S4),

$$\begin{aligned}
\hat{S}(\alpha) &= \exp(\ln \alpha \cdot \hat{g}) \\
&= \exp[(i\pi + \ln|\alpha|)\cdot \hat{g}] \\
&= \exp(i\pi \cdot \hat{g})\cdot \exp(\ln|\alpha|\cdot \hat{g}) \\
&= \hat{S}(-1)\cdot \hat{S}(|\alpha|), \;\; (\alpha < 0).
\end{aligned} \tag{S9}$$

The above result implies that a TS operator with negative factor $\alpha$ is equivalent to successive actions of temporal scaling with factor $|\alpha|$ followed by flipping the sequence over the time axis.

##### 1.3 Temporal scaling operators acting in succession

In the main text, we described that the successive action of two temporal scaling operators on an arbitrary sequence can be replaced by another operator, which writes:

$$\hat{S}(\alpha) \cdot \hat{S}(\beta) \cdot \mathbf{y}(t) = \hat{S}(\beta) \cdot \hat{S}(\alpha) \cdot \mathbf{y}(t) = \hat{S}(\alpha\beta) \cdot \mathbf{y}(t) = \mathbf{y}(\alpha\beta t). \tag{S10}$$

It is worth noting that this equation only holds when $\alpha$ and $\beta$ are both larger than 0. When two operators with opposite signs or with both negative signs are applied, it yields:

$$\hat{S}(\alpha) \cdot \hat{S}(\beta) \cdot \mathbf{y}(t) = \begin{cases} \mathbf{y}(t_\infty + \alpha\beta t) & \alpha \geq 0 \;\; \beta < 0, \\ \mathbf{y}(t_0 + (t_\infty - t_0)/\beta + \alpha\beta t) & \alpha < 0 \;\; \beta \geq 0, \\ \mathbf{y}(t_\infty - (t_0 - t_\infty)/\beta + \alpha\beta t) & \alpha < 0 \;\; \beta < 0, \end{cases} \tag{S11}$$

which are not considered in the present study.

##### 1.4 Temporal dynamics of scaled sequence and corresponding neural representation

Next we derive the temporal dynamics of consequential sequences after temporal scaling. To study the temporal dynamics, we apply a temporal derivative operator $\partial_t$ to the scaled sequence, and it yields:

$$\partial_t(\mathbf{y}(\mathbb{t}) = \partial_t[\hat{S}(\alpha) \cdot \mathbf{y}(t)] = \partial_t[\hat{S}(\alpha)] \cdot \mathbf{y}(t) = \Big[\sum_{n=0}^{\infty} \frac{(\ln\alpha)^n}{n!} \partial_t(t\partial_t)^n\Big] \cdot \mathbf{y}(t). \tag{S12}$$

To calculate the above equation, we should further expand $\partial_t(t\partial_t)^n$. By recursively applying $\partial_t$ on each $t\partial_t$ in $(t\partial_t)^n$, we derive:

$$\partial_t(t\partial_t)^n = (1 + t\partial_t)\partial_t(t\partial_t)^{n-1} = \cdots = (1 + t\partial_t)^n \partial_t. \tag{S13}$$

Substituting the above equation into Eq. S12, we can express the temporal dynamics of the scaled sequence as:

$$\begin{aligned} \partial_t \mathbf{y}(\mathbb{t}) &= \Big[\sum_{n=0}^{\infty} \frac{(\ln\alpha)^n}{n!} \partial_t(t\partial_t)^n\Big] \cdot \mathbf{y}(t) \\ &= \Big[\sum_{n=0}^{\infty} \frac{(\ln\alpha)^n}{n!} (1 + t\partial_t)^n \partial_t\Big] \cdot \mathbf{y}(t) \\ &= \exp[\ln\alpha(1 + t\partial_t)] \cdot \partial_t \mathbf{y}(t) \\ &= \alpha\hat{S}(\alpha) \cdot \partial_t \mathbf{y}(t) \end{aligned} \tag{S14}$$

Similar to the above analysis, the temporal scaled dynamics of neural responses can be derived as:

$$\begin{aligned} \partial_t u[z(\mathbb{t})] &= \Big[\sum_{n=0}^{\infty} \frac{(\ln\alpha)^n}{n!} \partial_t[t(\partial_t z)\partial_z]^n\Big] \cdot u[z(t)] \\ &= \Big[\sum_{n=0}^{\infty} \frac{(\ln\alpha)^n}{n!} (1 + [t(\partial_t z)\partial_z])^n \partial_t\Big] \cdot u[z(t)] \\ &= \exp\Big[\ln\alpha(1 + [t(\partial_t z)\partial_z])\Big] \cdot \partial_t u[z(t)] \\ &= \alpha\hat{S}_u(\alpha) \cdot \partial_t u[z(t)] \end{aligned} \tag{S15}$$

#### 2 Continuous attractor network dynamics

We described in the main text a neural circuit model named CAN, whose dynamics writes:

$$\tau\dot{u}(x,t) = -u(x,t) + \rho \int J(x,x')r(x',t)dx' + I_{ext}, \tag{S16a}$$

$$r(x,t) = \frac{[u(x,t)]_+^2}{1 + k\rho \int u[(x',t)]_+^2 dx'}. \tag{S16b}$$

Given that neurons in the circuit are connected reciprocally by a Gaussian-shaped function, we postulated that neural responses of the attractor states are expressed as follow:

$$\bar{u}(x,t) = A_u \exp[-(x-z(t))^2/4a^2], \quad \bar{r}(x,t) = A_r \exp[-(x-z(t))^2/2a^2], \tag{S17a}$$

where $\bar{u}(x,t)$ and $\bar{r}(x,t)$ are both Gaussian curves centered at the represented position $z(t)$.

#### 2.1 Perturbation analysis of Continuous Attractor Networks

A CAN only reserves those perturbations parallel to its $z$ manifold. To verify this point, we perform the perturbation analysis, and calculate the eigenvalue and eigenfunctions of the time derivative operator. The neural activity $u(x,t)$ of the CAN can be expressed as a combination of the attractor state and a perturbation: $u(x,t) = \bar{u}(x-z) + \delta u(x,t)$. Substituting it into the network dynamics (Eq. S16a), we derive the dynamics of the perturbation:

$$\tau \frac{d}{dt}\delta u(x,t) = -\delta u(x,t) + \int K(x,x'|z)\delta u(x',t)dx', \tag{S18}$$

where

$$K(x,x'|z) = \rho \int J(x-x'')\partial \bar{r}(x''-z)/\partial \bar{u}(x'-z)dx''. \tag{S19}$$

Based on the expression of $r(x,t)$ in Eq. (S16b), we can calculate $K(x,x'|z)$ as follow:

$$\begin{aligned}
&\int K(x,x'|z)\delta u(x',t)dx' \\
&= \int \Big[\frac{2\rho u(x',t)}{D} J(x,x')\delta u(x',t) - J(x,x')k\rho\frac{u^2(x',t)}{D^2}\int 2u(x'',t)\delta u(x'',t)dx''\Big]dx' \\
&= \int \frac{2\rho u(x',t)}{D} J(x,x')\delta u(x',t)dx' - \iint 2k\rho J(x,x')\frac{u^2(x',t)}{D^2} u(x'',t)\delta u(x'',t)dx'dx'' \\
&= \int \frac{2\rho u(x',t)}{D} J(x,x')\delta u(x',t)dx' - \iint 2k\rho J(x,x'')\frac{u^2(x'',t)}{D^2} u(x',t)\delta u(x',t)dx''dx' \\
&= \int \frac{2\rho u(x',t)}{D}\Big[J(x,x') - k\rho\int J(x,x'')\frac{u^2(x'',t)}{D}dx''\Big]\delta u(x',t)dx' \\
&= \int \frac{2\rho u(x',t)}{D}\Big[J(x,x') - k\rho\int J(x,x'')r(x'',t)dx''\Big]\delta u(x',t)dx'.
\end{aligned} \tag{S20}$$

Therefore, we conclude that

$$K(x,x'|z) = \frac{2\rho u(x',t)}{D}\Big[J(x,x') - k\rho\int J(x,x'')r(x'',t)dx''\Big], \tag{S21}$$

where $D$ denotes the nominator of $r(x,t)$, which can be calculated from $D = 1 + k\rho\int \bar{u}^2(x',t)dx'$.

Here, we treat $\int K(x,x'|z)(*)dx'$ as a linear operator, and consider it in a Hilbert $L^2$ space constructed by a set of basis functions,

$$v_n(x|z) = \frac{(-1)^n(\sqrt{2}a)^{n-1/2}}{\sqrt{\pi^{1/2}n!2^n}} \exp\Big[\frac{(x-z)^2}{4a^2}\Big]\Big(\frac{d}{dx}\Big)^n \exp\Big[\frac{(x-z)^2}{2a^2}\Big]. \tag{S22}$$

In this Hilbert space, we can calculate the eigenvalues and corresponding eigenfunctions of the operator $\int K(x,x'|z)(*)dx'$. We list the eigenvalues as follow,

$$\lambda_1 = 1, \quad \lambda_2 = 1 - \sqrt{1 - J_c^2/J_0^2}, \quad \lambda_n = 2^{2-n} \ (n \geq 3), \tag{S23}$$

where $J_c$ denotes the critical connection weight above which the network can hold a stable non-zero activity. Eq. (S23) shows that only perturbations parallel to the corresponding eigenfunction of $\lambda_1$ can be reserved by the network. This eigenfunction happens to be parallel to the translation direction along $z$ space,

$$f_1(x|z) = v_1(x|z) \propto (x-z)\exp\Big[\frac{(x-z)^2}{4a^2}\Big], \tag{S24}$$

which implies that the neural activity bump can move smoothly along $z$ direction.

#### 2.2 Calculation of bump amplitude and velocity

With projection method adopted from [2], we can theoretically calculate the magnitude and moving velocity of neural activities in the CAN. Substituting the neural responses (Eq.S17) into the network dynamics Eq. S16a, we obtain that:

$$\text{LHS} = \tau \frac{A_u}{2a^2}(x-z)\exp[-(x-z(t))^2/4a^2]\cdot\frac{dz}{dt}, \tag{S25a}$$

$$\text{RHS} = (-A_u + \sqrt{\pi}a\rho J_0 A_r)\exp[-(x-z(t))^2/4a^2] + \text{I}_0\exp[-(x-z_\infty)^2/4a^2]. \tag{S25b}$$

Substitute the solutions into Eq. S16b, then we obtain the relationship between $A_u$ and $A_r$:

$$A_r = \frac{A_u^2}{1+k\rho\sqrt{2\pi}aA_u^2} \tag{S26}$$

As stated in SI. Sec. 2.1, the CAN dynamics can be considered along a set of basis functions. The first two of them are shown as follow, which dominate the amplitude and spatial translation of the neural response respectively:

$$f_1(x|z) = (x-z)\cdot\exp[-(x-z(t))^2/4a^2], \tag{S27a}$$

$$f_2(x|z) = \exp[-(x-z(t))^2/4a^2]. \tag{S27b}$$

Then we can project Eq. S25 onto these basis functions to study its corresponding properties. By projecting function $f(x)$ onto $\phi(x)$, we mean that we compute their inner project that is defined as an integral $\int_x f(x)\phi(x)dx/\int_x \phi^2(x)dx$.

Projecting both sides of Eq. S25 onto $f_2$ and considering a weak input condition ($\text{I}_0 \approx 0$), we obtain that:

$$0 = (-A_u + \sqrt{\pi}a\rho J_0 A_r)\cdot\sqrt{2\pi}a. \tag{S28}$$

Given Eq. (S26 and S28), we can calculate the amplitude $A_u$ of neural response:

$$A_u = \frac{\sqrt{\pi}a\rho J_0 + \sqrt{\pi a^2\rho^2 J_0^2 - 4k\rho\sqrt{2\pi}a}}{2k\rho\sqrt{2\pi}a}. \tag{S29}$$

According to the above equation, $J_0$ has to be larger than a critical value $J_c$ to allow $A_u$ to be real. Equating the term under the root sign with 0, we obtain the expression of $J_c$:

$$J_c = \sqrt{\frac{4\sqrt{2\pi}k}{\pi a\rho}} \tag{S30}$$

Projecting Eq. S25 onto $f_1$ yields:

$$\text{LHS} = \tau a A_u\frac{\sqrt{2\pi}}{2}\cdot\frac{dz}{dt}, \tag{S31a}$$

$$\text{RHS} = 0 - a\text{I}_0\frac{\sqrt{2\pi}}{2}\cdot(z-z_\infty)\cdot\exp[-(z-z_\infty)^2/8a^2]. \tag{S31b}$$

Equating both sides, we derive a differential equation of the center of the neural response:

$$\frac{dz}{dt} = -\frac{\text{I}_0}{\tau A_u}\cdot(z-z_\infty)\cdot\exp[-(z-z_\infty)^2/8a^2]. \tag{S32}$$

We solve this equation by separating the variables and integrating both sides of it. Since the integral over $z$ is not analytically solvable, we expressed it by the exponential integration function $\mathsf{Ei}(x) = \int_{-\infty}^{x} e^t/t dt$:

$$\begin{aligned}\int_{t_1}^{t_2} dt &= \int_{z_1}^{z_2}(z-z_\infty)^{-1}\exp[(z-z_\infty)^2/8a^2]dz \\ &= \int_{z_1}^{z_2}\frac{1}{2}(z-z_\infty)^{-2}\exp[(z-z_\infty)^2/8a^2]dz^2 \\ &= \int_{(z_1-z_\infty)^2/8a^2}^{(z_2-z_\infty)^2/8a^2}\frac{1}{2}w^{-1}\exp(w)dw \\ &= \frac{1}{2}\big[\mathsf{Ei}((z_2-z_\infty)^2/8a^2) - \mathsf{Ei}((z_1-z_\infty)^2/8a^2)\big],\end{aligned} \tag{S33}$$

where $z_1, z_2$ represent 2 arbitrary states between $z_0$ and $z_\infty$, while $t_1, t_2$ represent their corresponding time step. Then the time duration of the sequence can be written as:

$$T = \frac{1}{2}\big[\mathsf{Ei}((z_\infty - z_\infty)^2/8a^2) - \mathsf{Ei}((z_0 - z_\infty)^2/8a^2)\big]. \tag{S34}$$

#### 3 Temporal scaling operator in the neural circuit dynamics

We perform theoretical analysis to verify the emergence of temporal scaling operator in the CAN circuit. The neural activity at an arbitrary time is represented as:

$$u(x,t) = \bar{u}(x, z(t)) + \delta u(x,t). \tag{S35}$$

Substituting it into the network dynamics and utilizing the basis functions we have defined in Sec. 2.1, we obtain that:

$$\begin{aligned} \tau\frac{du(x,t)}{dt} =& - [\bar{u}(x,z(t)) + \delta u(x,t)] + \rho \int J(x,x')\bar{r}(x',t)dx' \\ &+ \int K(x,x'|z)\delta u(x,t)dx' + \alpha \mathrm{I}_0(x|z_\infty), \\ =& \sum_n b_n(\lambda_n - 1)f_n(x|z) + \alpha\sum_n a_n f_n(x|z), \end{aligned} \tag{S36}$$

where we took advantage of the properties of the attractor states to cancel the two terms of $-\bar{u}(x,z(t))$ and $\rho\int J(x,x')\bar{r}(x',t)dx'$ with each other. Moreover, we decomposed $\delta u$ and $I$ into components along the basis functions,

$$\delta u(x,t) = \sum_n b_n f_n(x|z), \tag{S37a}$$

$$\mathrm{I}_0(x|z_\infty) = \sum_n a_n f_n(x|z). \tag{S37b}$$

As stated in SI. Sec. 2.1, only perturbations along $z$ direction can be preserved by the network dynamics, so we can expand $d/dt$ according to the chain rule and omit the partial derivatives respective to the variables except $z$:

$$\tau\frac{dz}{dt}\frac{\partial}{\partial z}u(x,t) = \sum_n b_n(\lambda_n - 1)f_n(x|z) + \alpha\sum_n a_n f_n(x|z). \tag{S38}$$

Since the basis functions $f_n(x|z)$ are orthogonal to each other, Eq. S38 can be considered as a set of independent equations. We derived that $\lambda_1 = 1$ in SI. Sec. 2.1, so the first term on the right hand side of Eq. S38 will vanish when $n = 1$. To simplify Eq. S38, we calculate the inner product of both sides of it with $f_1(x|z)$:

$$\tau\frac{dz}{dt}\Big\langle \partial_z(\sum_n c_n f_n(x|z))\Big| f_1(x|z)\Big\rangle = \sum_n b_n(\lambda_n - 1)\Big\langle f_n(x|z)\Big| f_1(x|z)\Big\rangle + \alpha\sum_n a_n\Big\langle f_n(x|z)\Big| f_1(x|z)\Big\rangle, \tag{S39}$$

where $\langle f(x)|g(x)\rangle$ denotes the inner product of $f(x)$ and $g(x)$, i.e., $\int f(x)g(x)dx$. Consider the orthogonality of $f_n(x|z)$ and the relationship of $\partial_z f_2(x|z) \propto f_1(x|z)$, we obtain that,

$$\tau c_2\frac{dz}{dt} = \alpha a_1, \tag{S40}$$

where $c_2$ is the second component of $u(x,t)$, representing the bump amplitude of the neural activity, and $a_1$ is the first component of $I(x|z_\infty)$. Therefore, we can substitute $c_2 = A_u$ and $a_1 = \Big\langle I(x|z_\infty)\Big| f_1(x|z)\Big\rangle$ into Eq. (S40) and it yields,

$$\tau A_u\frac{dz}{dt} = \alpha \mathrm{I}_0(z_\infty - z)\exp[-(z_\infty - z)^2/8a^2]. \tag{S41}$$

We see from Eq. S41 that the gain factor $\alpha$ and spatial offset $z_\infty - z$ constitute the temporal scaling operator.

#### 4 Simulation details

##### 4.1 The simulation of the CAN

We simulated a continuous attractor network with 512 neurons. The preferred positions of these neurons are set uniformly distributed in the $z$ manifold. Without loss of generality, we set the $z$ manifold spans in the range of $[-\pi, \pi)$ as a dimensionless number. To avoid the boundary effect, we applied a periodic condition in our simulations by connecting the $-\pi$ and $\pi$ of the $z$ manifold. Since the range of $[-\pi, \pi)$ is much larger than the connection width between neurons ($a = 0.5$, Eq. 14), the calculations where integrals are calculated over an infinite range are not substantially affected. The differential equations are simulated by Euler method and the time step $dt$ in simulation is set to be $1/10$ of time constant $\tau$. Other parameters are listed in Table. S1. The network was coded in Python and run on a MacBook Pro laptop with M1Pro CPU and 32GB RAM.

Table S1: Parameters of CAN simulations

| Symbol | Description | Value |
|---|---|---|
| $N$ | Number of neurons | 512 |
| $\rho$ | Neuron density | $N/2\pi$ |
| $a$ | Tuning width | 0.5 |
| $k_c$ | Critical inhibitory strength | 12.766 |
| $k$ | Inhibitory strength | $0.04k_c$ |
| $\tau$ | Time constant of neural activities | 1ms |
| $dt$ | Time step in numerical simulations | $0.1\tau$ |
| $J_0$ | Connection strength | 1 |
| $z_0$ | Position of the initial state | $2\pi/5$ |
| $z_\infty$ | Position of the target state | 0 |
| $\mathrm{I}_0$ | Referenced input strength | 0.04 |
| F | Fano factor of Poisson noise | 0.001 |

##### 4.2 Dimension reduction

To reduce the dimensions of the neural activities, we applied principle component analysis (PCA) to the network data. For each scaling factor $\alpha$, we ran the network for 10 times to obtain a $10 \times T \times N$ neural activity matrix ($T$ for number of time steps and $N$ for number of neurons). The activities when $\alpha = 1$ and $T = 1$ were taken to build the projection vectors and 3 PCs were preserved to explain the variance over 10 noisy samples. Therefore, the dimension of the network data is reduced from $N$ to 3. With the projection vectors we have built, the data of each $\alpha$ value and each time step were projected into the 3 dimenisonal space and formed the low-dimensional manifold depicted in the main text Fig. 3C. Note that 3 PCs are sufficient to explain most of the variance of the neural activities, as shown in Fig. S1.

##### 4.3 The simulation and training of handwritten digits

To verify the neural sequence generated by the CAN in the disentangled circuit can produce complex sequence with different scales, we trained a feedforward network that uses the activity of CAN to generate the trajectory of the handwritten digit "6".

We constructed a simple feedforward network consisting of three-layer of neurons with 100, 20, and 2 neurons in each layer, respectively. The input of the feedforward network is the spatiotemporal responses of the 512 neurons in the CAN. And the first and the second layers in the feedforward network use ReLU and tanh as activation functions respectively, in order to best fit the shape of handwritten digits. The neurons in final layer is linear and directly outputs the coordinates $(x, y)$ of the digit's trajectory. The handwritten digit trajectory data was obtained from [3].

Consistent with the previous simulation, we used CAN with referenced input strength $I_0$ to train the three-layer network. The target $(x_{target}(t), y_{target}(t))$ at each time step $t$ was calculated based on the position of the CAN bump center. During the training process, we fed the CAN sequence with random noise ($\sim \mathcal{N}(0, 0.001)$) step by step. The error was calculated by

$$Err = \sum_{t=0} \frac{1}{2}[(x(t) - x_{target}(t))^2 + (y(t) - y_{target}(t))^2].$$

We used the adaptive moment estimation stochastic gradient descent algorithm (Adam) implemented in Pytorch to minimize the mean square error (MSE) between the network output and the target trajectory coordinates. The learning rate was set to 0.002, and all other parameters were set to Pytorch defaults. The network was trained for a total of 400 epochs before stopping.

We trained digits "0-9". The training loss for number "6" is shown in Fig. S2. The results of all the other numbers are shown in Fig. S3. Please note that the color bars used for the target and digital trajectories in the main text and supplementary are different for visual purposes. However, the target and $\alpha = 1$ traces are at about the same time.

#### 5 Supplementary Figures

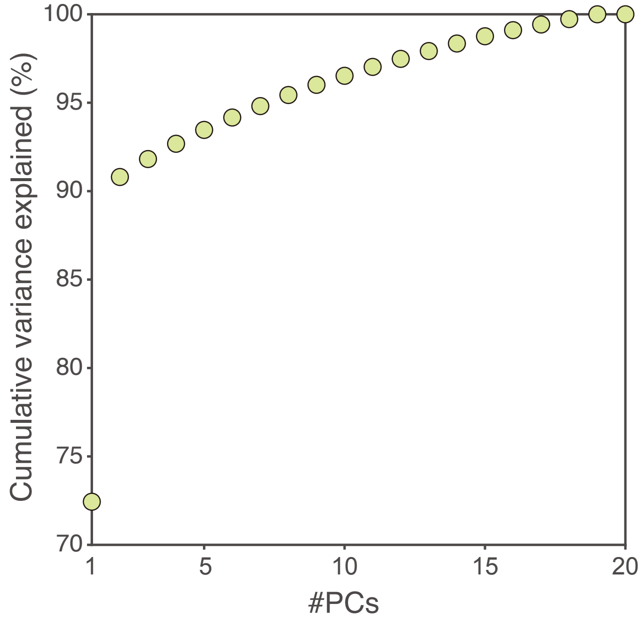

Figure S1: Cumulative variance explained by number of PCs

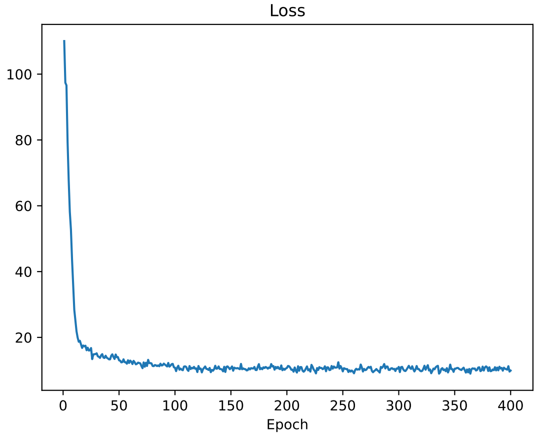

Figure S2: The loss of training digital number "6" in the main text.

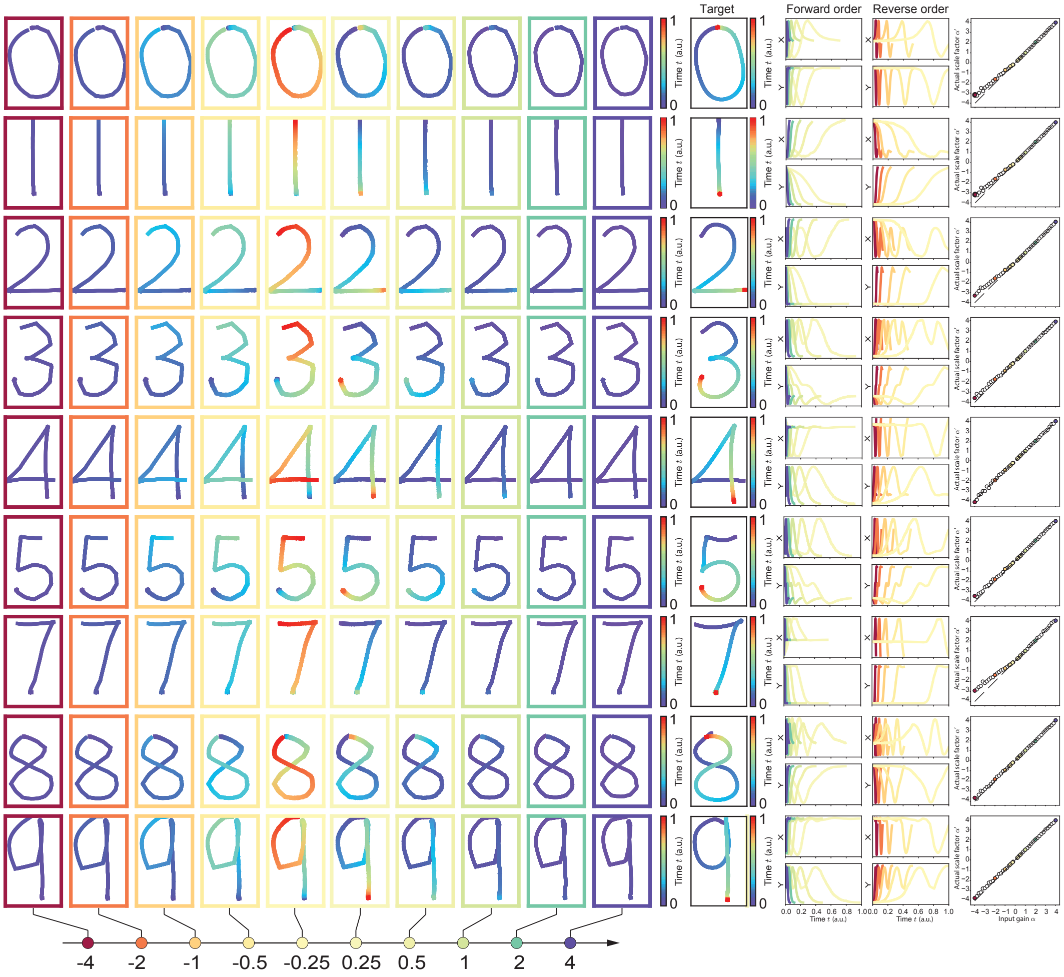

Figure S3: Results of training for digit "0-9" except "6"
